# Mean environmental associations obscure drivers of zooplankton community dynamics

**DOI:** 10.64898/2026.04.16.718861

**Authors:** Miriam Beck, Louise Laux, Jean-Olivier Irisson, Franziska Schrodt, Luca Santini

## Abstract

Zooplankton communities are influenced by multiple environmental factors, including temperature, nutrient and resource availability, which fluctuate seasonally and across years. While long-term average effects can identify overall drivers, they may overlook dynamic, context-dependent effects that govern short-term changes in diversity and abundance. Understanding and disentangling both perspectives is crucial for identifying and estimating the drivers that shape community structure under varying environmental states.

Here, we applied Empirical Dynamic Modeling (CCM, SMap) to a 12-year weekly zooplankton time series to identify causal environmental drivers of taxonomic and morphological diversity and quantify how the influence of each driver shifts over time. We contrast these results with static long-term average effects inferred from Generalized Linear Models which included predictor sets identified using covariate adjustment and accounting for temporal autocorrelation.

Drivers linked to long-term average associations differed from those regulating short-term zooplankton dynamics, revealing a decoupling between mean environmental effects and the drivers of temporal variability. Temperature emerged as a persistent regulator of zooplankton dynamics across multiple diversity dimensions, while variables commonly associated with background trophic conditions (*e.g*. particulate organic matter) were primarily associated with long-term patterns and showed limited dynamical relevance. Importantly, we find evidence for morphological homogenisation in response to short-term fluctuations in chlorophyll *a*, which was not detectable in long-term average relationships.

This contrast highlights that mean environmental associations do not necessarily reflect the mechanisms governing community dynamics. Impacts might be underestimated if average effects appear weak, or misinterpreted if arising mainly from shared trends or seasonality rather than direct mechanisms Integrating both perspectives clarifies the identity and role of environmental drivers, improving inference and prediction of zooplankton community change through time.

## Introduction

Zooplankton communities are shaped by a suite of environmental variables that interact across temporal scales: Temperature directly constrains metabolic rates and phenology (Gillooly 2000, Schlüter et al. 2010, Svendsen et al. 2025) and directly and indirectly governs resource availability (Lewandowska et al. 2012). Nutrient supply regulates energy supply through bottom-up pathways (Sterner & Elser 2002) and physical processes such as water column mixing further mediate community responses of phyto- and zooplankton (Berline et al. 2012, Gonzalez-Gil et al. 2015, Wang et al. 2024). These drivers fluctuate seasonally (Record et al. 2010, Sommer et al. 2012, Wang et al. 2024) and interannually (Record et al. 2010, Gabaldón et al. 2019), and often exhibit long-term trends (Record et al. 2010, Beck et al. 2023). In consequence, their effects on community structure are often nonlinear, lagged, or depend on community/environmental state (Obertegger et al. 2007, Sinclair et al. 2021) or the temporal scale considered (Record et al. 2010, Gabaldón et al. 2019). Such temporal complexity challenges the identification of environmental drivers from time series data even when expected underlying causal relationships are conceptually well understood (Fig. 1).

**Fig. 1:**
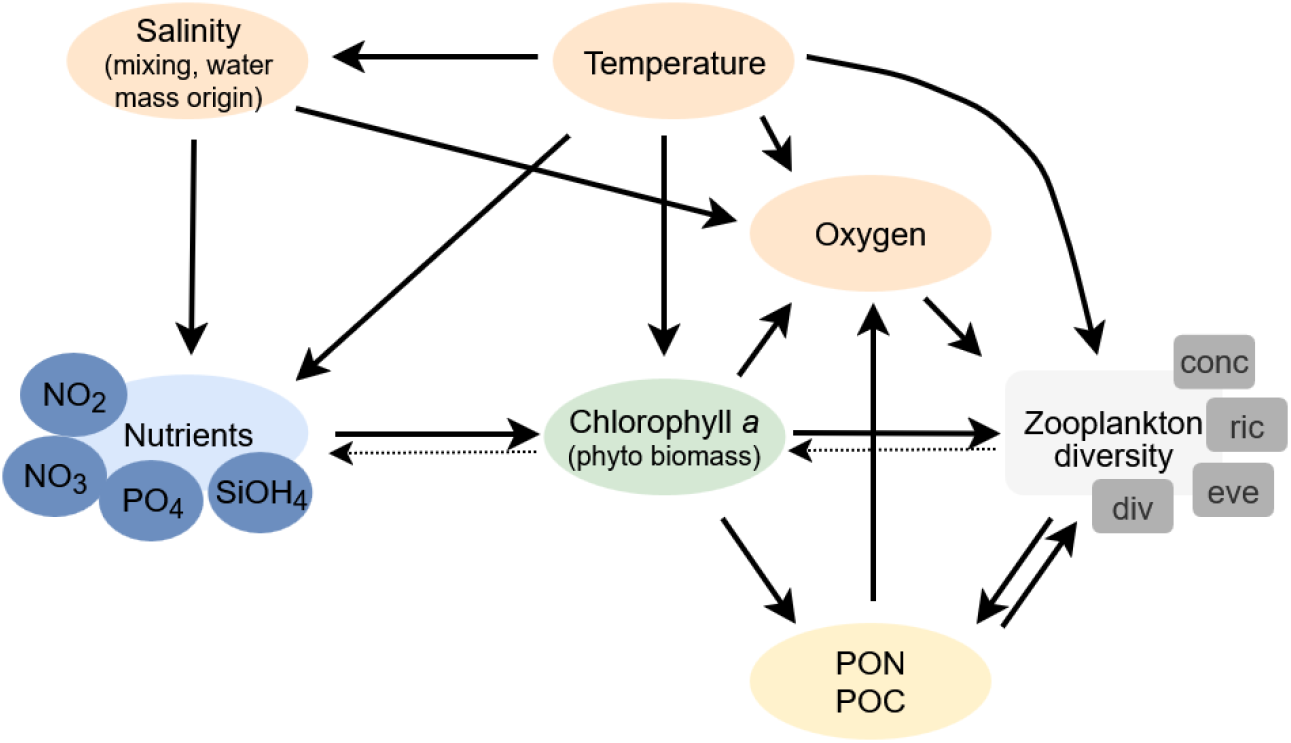
Causal model of expected/known relationships between environmental variables and zooplankton diversity. Nodes represent environmental variables included in the analysis. Arrows indicate expected causal relationships: a change in the upstream node is expected to induce a change in the downstream node either directly or indirectly via the indicated unobserved mediating process). Dashed arrows indicate known feedback effects. Some variables represent proxies for underlying processes (e.g. salinity) or ecological state (e.g. chlorophyll a).

This challenge is particularly pronounced for diversity, which reflects combined demographic responses, shifts in relative abundance, and species turnover of multiple taxa, thus integrating processes operating on different temporal scales. Albeit individual species responding predictably to environmental drivers, diversity can exhibit nonlinear and context-dependent behaviour, where its response depends on the current environmental conditions and community composition rather than any single variable consistently through time. Consequently, statistical associations between environmental variables and diversity may obscure the dynamical processes governing community change.

Generally, standard regression approaches are vulnerable to collinearity, confounding, and spurious associations if relying on observational data (Arif & MacNeil 2022): A variable may appear associated with diversity not because it causally regulates community dynamics, but because both respond to the same seasonal cycle or long-term trend. Moreover, average statistical associations often fail to capture the nonlinear or context-dependent processes governing community change at shorter timescales (Hastings et al. 2018), causing variables that strongly influence fluctuations to appear weak or irrelevant in long-term associations. In contrast, long-term associations between environmental factors and zooplankton diversity can successfully inform prediction or forecasting tasks (Villarino et al. 2015) but they may overestimate the role of background factors and underestimate drivers of week-to-week variability. Thus, distinguishing statistical association from dynamical causation remains a central challenge in ecology with important implications for management and conservation. For example, if nutrient concentrations are associated with declining diversity only because both respond to warming or internal trophic feedback, reducing nutrient input alone would not reverse the decline.

While reported associations are often assessed relying on average changes and linear relationship (Mackas & Beaugrand 2010), their integration with explicitly dynamic frameworks, such as Empirical Dynamic Modelling (EDM) still is limited in ecology. EDM (Sugihara et al. 1997) relies on state-space reconstruction of time series, providing a toolbox to identify causal relationships between pairs of variables (via Convergent Cross-Mapping (CCM), Sugihara et al. 2012) and quantify their effect through time (via Sequential Locally Weighted Global Linear Maps (SMap), Sugihara et al. 2012). Unlike GLMs, which estimate average associations and might be influenced by extreme or anomalous periods, CCM identifies variables whose dynamics are causally embedded in the temporal change of zooplankton diversity (Fig. 2). Here, causal influence in the sense of dynamic systems means that the driver is part of the state-dependent rules governing how a system evolves; this effect can be non-linear, lagged, intermittent or masked by averages.

**Fig. 2:**
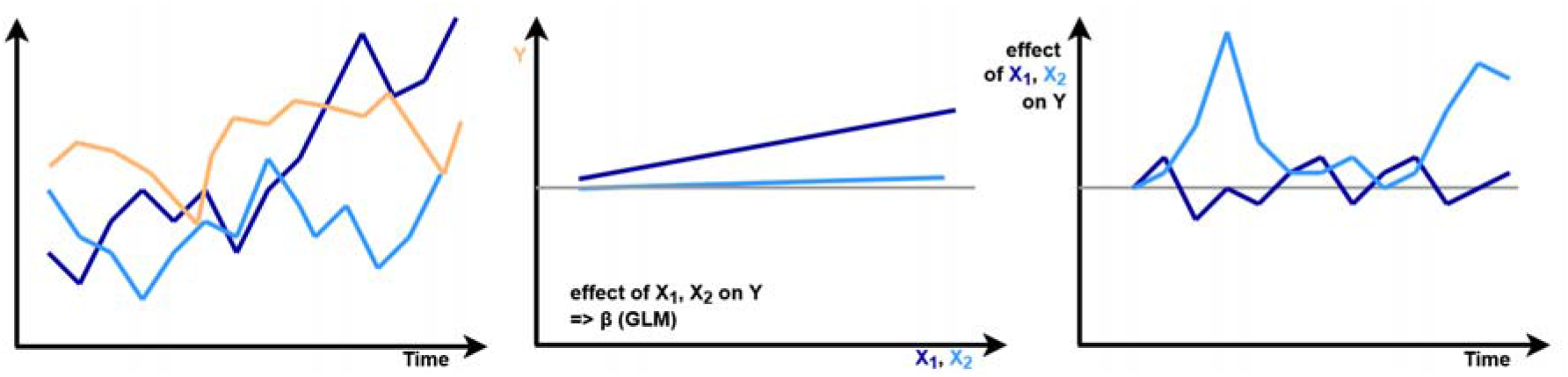
Visualisation of how the GLM (center) and SMap (right) quantify the effects of two driver time series (X_1_, X_2_, blue) on a response (Y, orange). While GLM identifies an overall small but positive effect of X_1_ on Y, its effect on Y’s dynamics is neglectable compared to that of X_2_. For this relationship, time-explicit effects capture the context-dependency of this effect which is strongest if values of X_2_ are low. CCM likely would identify X_2_, but not X_1_ as a driver of Y’s dynamics.

**Fig. 2:**
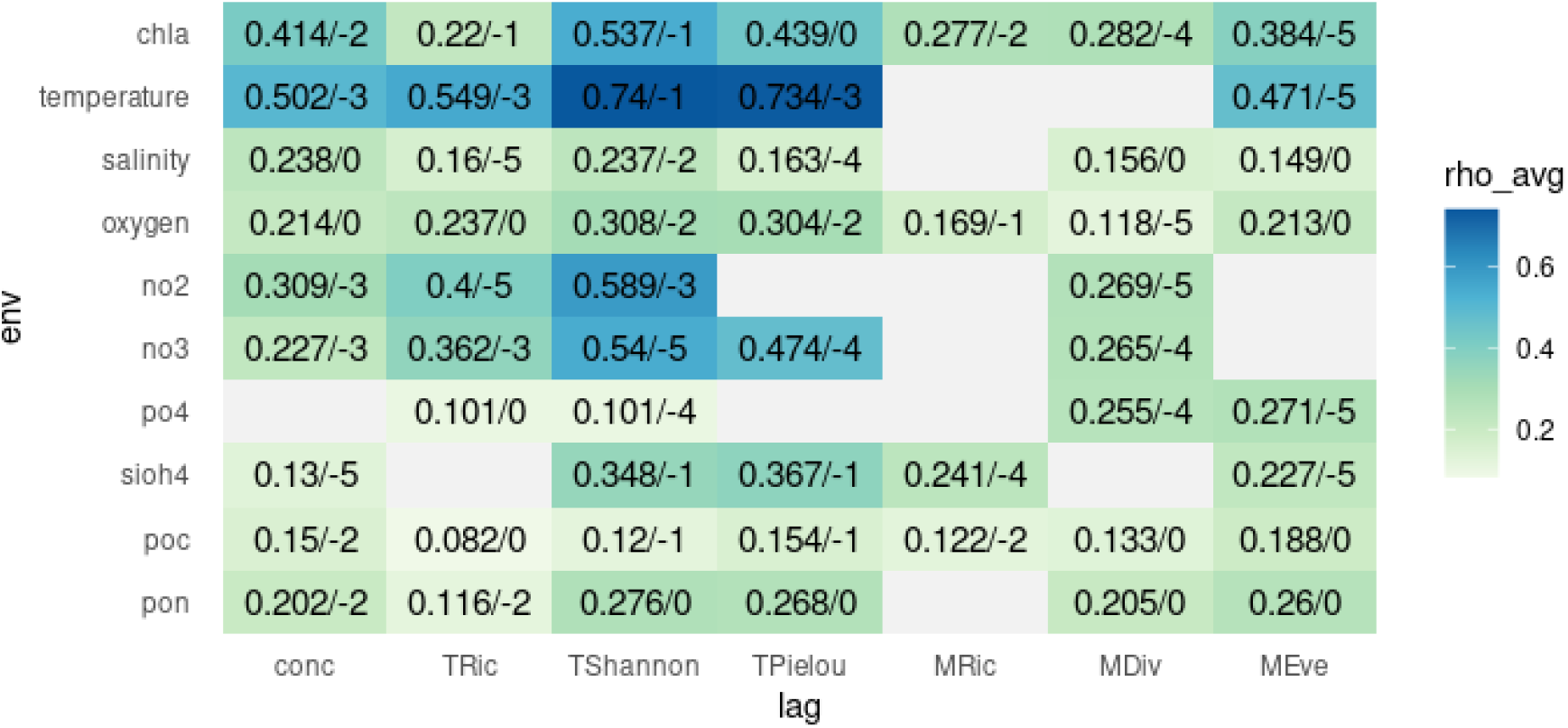
Pairwise relationships between environment variables as drivers of diversity dynamics at time lag showing the strongest prediction skill (0 to 5 week). The first labelling value indicates rho value, a measure of strength of predictability (which can be correlated to, but is not a measure of effect size), the second value indicates the time lag in weeks at which prediction skill was largest.

In plankton systems, this approach has successfully helped disentangling bottom-up form top-down driving in plankton system (Frossard et al. 2018), or the drivers of phytoplankton populations (Mondal et al. 2022) or communities under different nutrient regimes (Chang et al. 2022). Here, we rely on its complementary view on ‘drivers’ to better disentangle the nature of environment-community diversity relationships.

Using twelve years of weekly zooplankton community data (Beck et al. 2023) this study aims to identify the environmental drivers of taxonomic and morphological diversity dynamics using nonlinear, state-dependent causal analysis within the empirical dynamic modelling (EDM) framework. We contrast the drivers inferred from EDM with long-term average environmental effects estimated using GLMs.

We expect CCM to preferentially identify proximate drivers such as temperature and chlorophyll *a* (representing phytoplankton concentration) as drivers on zooplankton dynamics, while indirect drivers such as nutrient concentrations are expected to exhibit weaker effects or show up at larger time lags. In contrast, GLMs which identify drivers with persistent and directionally consistent effects, might obscure state-dependent variables but identify temperature effects under long-term warming at the site. Agreement between both approaches is expected for dominant, state-independent drivers (*i.e*. temperature), whereas discrepancies will indicate drivers whose effects vary in strength or sign depending on ecological context. As for environmental variables we expect diversity indices to be sensitive at time lags of 1 to 5 weeks depending on the underlying population dynamics. Because different processes underlie changes in community structure, we expect zooplankton concentration to respond at shorter time lags, diversity and evenness at short-medium lags (reflecting shifts in relative abundance) and taxonomic richness to respond at longer lags corresponding to species appearance/disappearance.

## Material and methods

### Data

The study site is situated in the north-western Mediterranean Sea at the entrance of the bay of Villefrance-sur-mer, France. Zooplankton and environmental variables (Chl *a*, temperature, salinity, NO_2_, NO_3_, PO_4_, SiOH_4_, POC, PON) cover weekly samples between July 2009 to December 2020. Zooplankton has been hauled vertically from 75m to the surface with WP2 net (200um mesh size), fixed with formalin and a representative fraction was digitalised using the ZoosSan (). Imaged individuals were separated and taxonomy identified based on 45 morphological features (*e.g*. area, perimeter, greyness). Taxonomic identification was verified by a human specialist. For each sampling day, taxonomic diversity (richness (=TRic), Pielou evenness (TPielou), Shannon diversity (TShannon) was calculated in addition to zooplankton concentration. In addition, the morphological features were used to classify all individuals based on their morphological features into 200 morphological groups using dimensional reduction and kMeans clustering. Based on these groups, morphological diversity metrics equivalent to the taxonomic ones (richness (=MRic), evenness (=MEve), divergence (=MDiv)) were computed following Villeger et al. (2008) for each sampling date.

Diversity and environmental time series were subjected to seasonal-trend decomposition by Loess (STL) enabling to to extract and subtract the long-term or seasonal signal, respectively. Details on sampling, measurements of environmental data and data processing are provided by Beck et al. (2023).

### Drivers of zooplankton dynamics

Pairwise-causal links between environmental variables and each diversity metric were identified using Convergent Cross-Mapping (CCM, Sugihara et al. 2012). CCM is a model-free approach rooted in non-linear dynamic systems theory that uses state-space reconstruction to detect causal links in deterministic systems. Causality is inferred relying on Takens’ theorem which implies that when two variables are dynamically coupled, the time series of one variable contains information about the underlying system dynamics driving the other.

In practice, CCM quantifies how well historical values of one variable (‘X’) can be reconstructed (‘cross-mapped’) from the time-delay embedding of another variable (‘Y’). If X causally influences Y (X → Y), values of X can be reliably predicted from the reconstructed state space of Y. Causality is inferred based on convergence: a causal link is supported when the correlation between observed and cross-mapped values increases with time series length, while low correlation or absence of convergence indicates no dynamic coupling. Via pair-wise testing, CCM further allows inference of the direction of causal link (X → Y, Y → X) or bidirectional interactions (X <-> Y).

Parameters were set as follows: The optimal embedding dimension (‘E’) for each variable was identified as the embedding of the first maximum of prediction strength over a range from 0-10. For each pair of environment-diversity variables CCM was used to test for a causal link of environmental variables on diversity using the optimal embedding of predictor variables (‘div’) and n=200 bootstrapping runs. CCM was run across varying time lags (tp=0 to -5, corresponding to lags of 0 to 5 weeks) to identify the time lag of strongest driver effect on diversity. The library size was set to time series length and by default cross-map skill was evaluated using leave-one-out cross-validation. All other parameters were used with default configuration.

The mean prediction skill ρ (‘rho’), *i.e*. the correlation between observed and cross-mapped values, was calculated across bootstrapping for each step library size. It measures how well the prediction matches reality but does not imply the magnitude/strength of effect.

The causality criterion of increasing prediction skill with increasing library size was tested in two ways: 1) significance and positive tau value of Kendall test and 2) that prediction skill at 20% maximum library size was significantly larger than zero (one-sided t-test).

To make sure that detected links were not spurious correlations driven by shared seasonal patterns, we applied a seasonal surrogate approach (Tsonis et al. 2015). Seasonal surrogate data were generated by first estimating the mean seasonal pattern (period = 52 weeks) and decomposing each time series into seasonal and residual components. The residuals of the target variable (‘env’) were then resampled, while preserving their autocorrelation structure, and recombined with the seasonal signal. CCM was then applied to each surrogate time series using the same model parameters, and this procedure was repeated 100 times to construct a null distribution of CCM skill (rEDMs ‘SurrogateData()’-function). If the initial rho value is significantly higher than the surrogates rho, we conclude the identified link to go beyond shared seasonal forcing.

All time series (environment, diversity) were scaled prior to running the analysis (Wang et al. 2020).

### Quantification of causal drivers

For each diversity metric, the set of identified causal drivers were then quantified across time using multi-variate Sequential Locally Weighted Global Linear Maps (S-Map). S-Map is a nonlinear, state-dependent extension of linear regression that estimates local interaction strengths by fitting weighted linear models in reconstructed state space, with greater weight given to nearby system states. This allows model coefficients to vary over time, thereby capturing possible temporal changes in the strength and direction of environmental effects on zooplankton diversity.

As predictors, we considered all environmental variables identified as significant drivers of the system in CCM at their optimal time lag (*i.e*. resulting in highest prediction skill; Wang et al. 2020). Considering the previously estimated embedding dimension, we then included the top E-1 predictors based on their prediction skill to avoid overparameterisation (Wang et al. 2020, Rigal et al. 2023). Theta (θ), describing the linearity of the system and governing a local *vs*. global forcing of predictors, was determined by evaluating prediction skill over a range of values (default; 0.01, 0.1, 0.3, 0.5, 0.75, 1, 1.5, 2, 3, 4, 5, 6, 7, 8, 9). We selected the theta corresponding to the first maximum of prediction skill. All further parameters remained unchanged from the default setting.

The resulting SMap coefficients for each predictor through time were summarised into average (mean, mean(abs)), variability (sd) and proportion of positive effects. CCM and SMap analysis was run using the rEDM package.

### Average long-term effects

To quantify the long-term average effect of environmental variables on zooplankton diversity, we fitted separate Generalized Linear Models (GLMs) for each diversity-environment pair. For each driver variable, we identified the necessary set of covariates based on known causal structure of the system (Fig.1, Tab. 1). Following the backdoor criteria from causal inference (*e.g*. Arif & MacNeil 2023, Schrodt et al. 2025), potential confounders - variables that influence both the exposure (environmental driver) and the response (diversity metric) - were included as covariates to block spurious associations, while avoiding adjustment for mediators or colliders to prevent bias (*i.e*. underestimation of effect in case of mediators; creating false association in case of colliders). Such ‘covariate adjustment’ ensures that estimated coefficients reflect the total effect of each environmental variable on zooplankton diversity, accounting for indirect pathways and shared drivers in the system. Models accounted for temporal autocorrelation using autoregressive structure (AR(1) or (AR(2)). The AR order was selected based on residual diagnostics and model fit (AIC).

**Table 1:** Covariate sets for estimating effects of each environmental variable. Example model: Div ∼ chl. a + temperature + salinity.

| Environmental driver | Adjusted covariate set |
| --- | --- |
| chl *a* | temperature, salinity |
| temperature | - |
| oxygen | chl. *a*, temperature, PON, POC |
| salinity | temperature, salinity |
| NO<sub>2</sub> | temperature, salinity |
| NO<sub>3</sub> | temperature, salinity |
| PO<sub>4</sub> | temperature, salinity |
| SiOH<sub>4</sub> | temperature, salinity |
| POC | chl. *a* |
| PON | chl. *a* |

The models were applied on de-seasonalised time series, removing a constant seasonal component from raw time series (see Beck et al. 2023) with scaled predictor variables.

## Results

### Causal drivers of diversity dynamics

For each diversity metric we identified four to ten significant drivers beyond the shared seasonal signal, with the prediction skills peaking at different time lags (0 to 5 weeks) (Fig. 2).

Temperature showed the highest prediction skill among all diversity variables, driving all metrics but morphological richness and morphological divergence. The optimal time lag varied between one and three weeks, but temperature appeared as the strongest predictor across time lags. Chlorophyll *a* and/or nitrogen concentration (NO_2_, NO_3_) also showed quite high predictability, notably for zooplankton concentration and taxonomic diversity indices. Despite consistently lower prediction skill relative to other environmental drivers, salinity, oxygen, and particulate organic matter (PON, POC) were identified as causal drivers for all or all but one diversity metrics. Phosphate and SiOH_4_ concentrations generally exhibited low prediction skill across environmental-diversity relationships, with exceptions for MDiv and MEve (phosphate) and TShannon and TPielou (SiOH_4_); however, even in these cases, predictability remained intermediate relative to other drivers. Generally, predictability of environment-diversity relationships were stronger for taxonomic metrics than for morphological ones, notably for physicochemical variables and nitrogen.

While some environmental variables were identified as causal predictors of a diversity variable across the whole range of time lags (*e.g*. concentration:temperature, TShannon: salinity/oxygen/no2), others only drove diversity dynamics at specific time lags (S2). Across diversity variables, the prediction skill of nutrient concentrations (NO_2_, NO_3_, PO_4_) increased with increasing time lag, while that of salinity and oxygen tended to decrease. Chlorophyll *a* generally showed highest prediction skill at time lags of one to two weeks, except for morphological divergence and morphological evenness where prediction was best at time lags of four and five weeks, respectively.

### Drivers of diversity dynamics through time

Multivariate SMaps included two to four (MDiv) predictors which had shown strongest predictive skill in CCM. Across diversity metrics, temperature clearly exerted the strongest effect, but at the same time the largest variability across time (Fig. 3a, b). While its effect was predominantly positive on diversity indices, its effect on concentration was exclusively negative (Fig. 3c). It was not included in the models of MDiv and MRic.

**Fig. 3:**
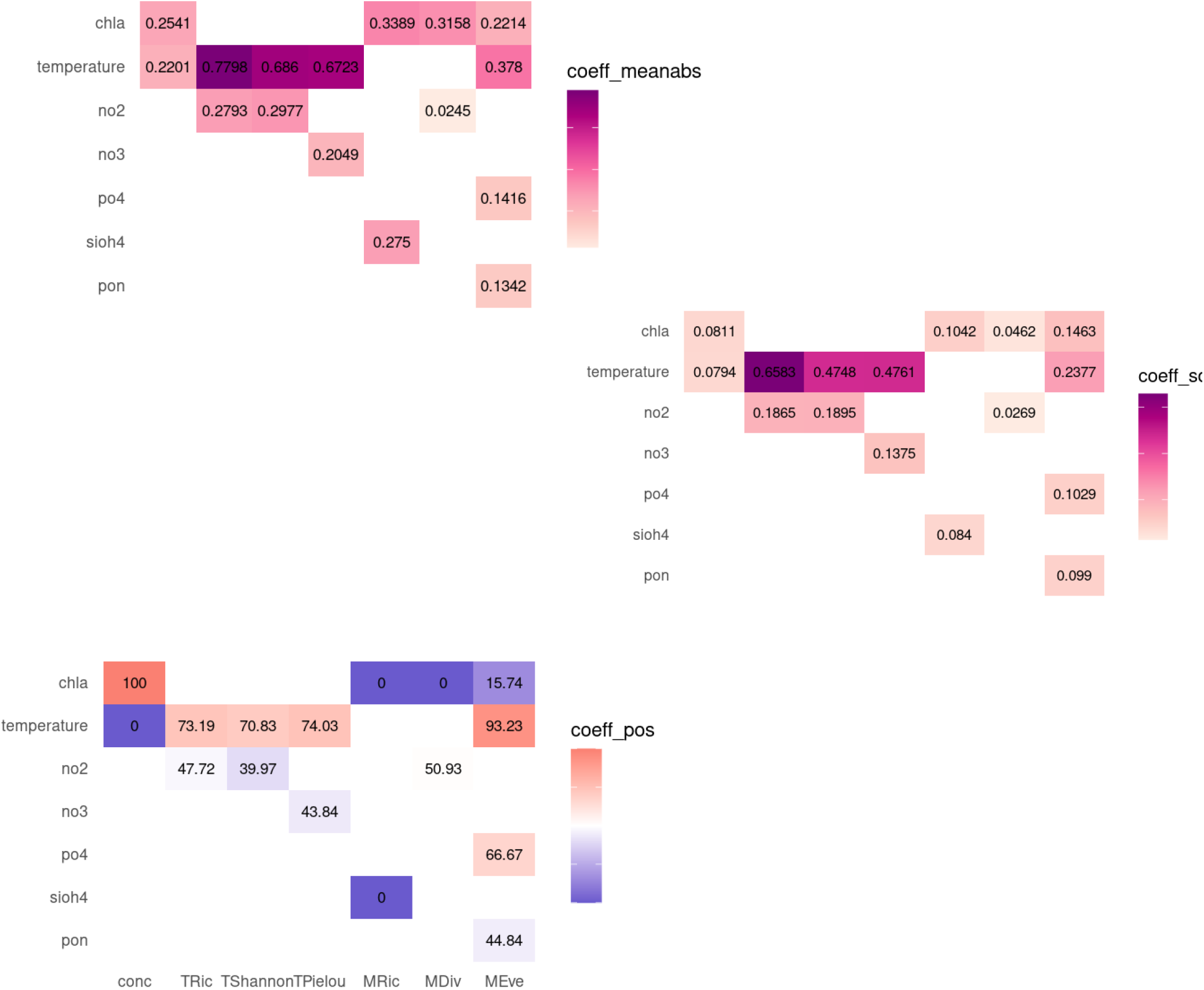
Average absolute effect strength (‘abs-mean’, A), effect variability (‘SD’, B) and proportion of positive effects (C) of environmental drivers on diversity. SMap coefficients were obtained from multivariate SMap using the predictors with highest prediction in CCM, and summarised across the whole 12-year time series.

**Fig. 4:**
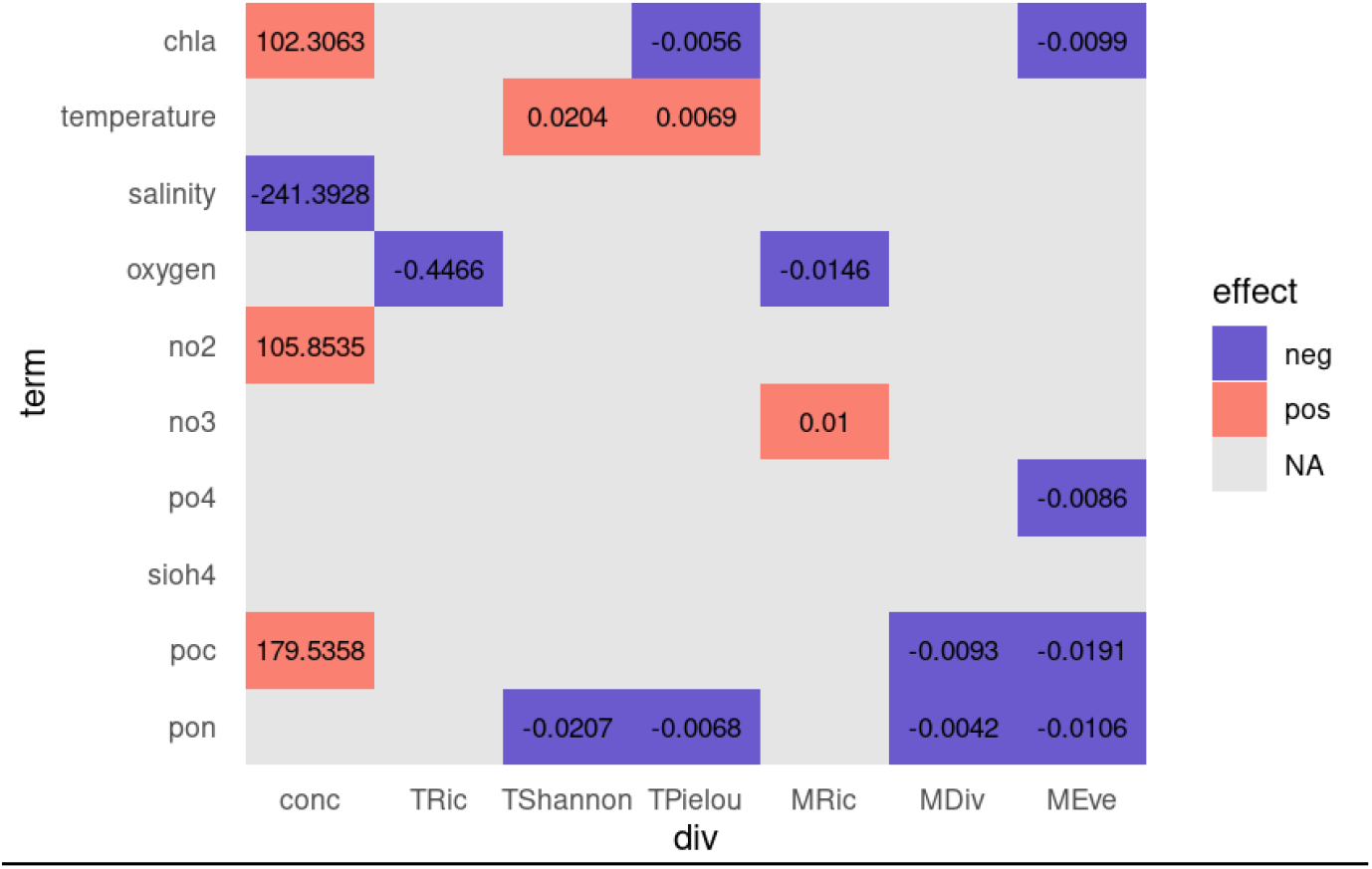
Average effects of environmental variables on diversity metrics. Effects were estimated as total effects comprising direct as well as indirect effects, while integrating confounding variables. Red and blue indicate significant positive and negative effect, respectively; lightgray the absence of significant effect.

The effects of NO_2_ and NO_3_ on taxonomic diversity (TRic, TShannon, TPielou) were smaller in absolute terms. Despite low variability in effect magnitude (SD), coefficients alternated between positive and negative values with negative effects predominating. NO_2_ effect on MEve was close to zero. Chlorophyll *a* was the strongest driver of dynamics of concentrations, MRic and MDiv, exhibiting very stable effects that were exclusively positive for concentrations and exclusively negative for MRic and MDiv. Silicate exhibited an intermediate, exclusively negative effect on MRic, while the effects of PO_4_ and PON on MEve were small compared to the stronger influences of temperature and chlorophyll *a*.

Model rho values ranged from 0.32 (MDiv) to 0.78 (TShannon), indicating a certain degree of predictability of zooplankton community diversity dynamics. As indicated by values of theta, the systems range from rather linear (θ=0.75-1.5: concentration, MDiv, MRic) to increasing non-linearity and strong state-dependency of environmental drivers (θ =3-4: TShannon/TPielou/MEve/TRic).

### Average long-term effect

GLMs identified one (TRic) to four (concentration, MEve) significant environmental drivers of zooplankton diversity in the long-term, which varied among diversity variables. Temperature increased taxonomic diversity and evenness, but did not significantly affect concentration, richness or morphological diversity. Higher chlorophyll *a* increased concentration, but reduced taxonomic and morphological evenness. Salinity and oxygen negatively impacted concentration and richness (taxonomic, morphological), respectively. Among nutrients, increasing nitrogen concentrations were linked to increasing concentration (NO_2_) and MRic (NO_3_), while increasing PO4 was associated with declining MEve. Particulate organic carbon was positively associated with zooplankton concentration but negatively affected MDiv and MEve; whereas particulate organic nitrogen was associated with declines in both taxonomic and morphorological diversity and evenness.

## Discussion

Using high-resolution long-term data of a plankton time series, we identified environmental drivers of community diversity dynamics as well as average associations. Although many identified drivers align with established zooplankton ecology, our analyses reveal that the mechanisms governing short-term community dynamics substantially differ from those shaping long-term averages. By disentangling state-dependent from mean effects across multiple diversity dimensions we demonstrate that commonly inferred environmental controls may obscure the mechanisms structuring temporal biodiversity fluctuations.

Among the environmental variables considered, temperature emerged as the only driver that consistently structured zooplankton dynamics across diversity metrics, particularly influencing community structure. This effect was rather stable across different time lags (S2), suggesting temperature as a continuous pervasive directly acting driver. Interestingly, average long-term associations were only detected for two taxonomic diversity metrics (TShannon, TEve), albeit it is often assumed a major long-term driver. In our analysis we excluded seasonality, suggesting that the often reported temperature effect is mainly due to seasonal coupling, which matches an observation in coastal plankton abundances (Wells et al. 2021). Although temperature effect on zooplankton dynamics was strongly structured by seasonality (S5), its effect goes beyond pure shared seasonal forcing as revealed by seasonal surrogate null models. The identification of temperature as a driver of various aspects of zooplankton dynamics suggests that ongoing warming will have consistent effects on community dynamics, even when long-term average relationships appear weak.

Chlorophyll *a* primarily acted as a driver of zooplankton biomass and exerted an exclusively positive effect on concentration dynamics. This was also reflected in the long-term average association, providing evidence for bottom-up support. In contrast, its influence on taxonomic diversity was comparatively weak across all-predictor SMaps (S4), supporting its role as a mediator variable primarily regulates zooplankton biomass (Basu and Pick 1997) rather than taxonomic community reorganization (Kâ et al. 2006, Picapedra et al. 2020, Weldrick et al. 2024). In contrast to taxonomic diversity, it was a major structuring driver for morphological diversity dynamics. Its effect was almost exclusively negative, supporting the hypothesis of functional (morphological) homogenisation under high-resource conditions, where increases in phytoplankton biomass favor a limited set of morphological traits and reduce trait variability without altering the overall species pool. Importantly, this effect on morphological diversity was mainly detectable through dynamic, state-dependent analyses and was not apparent in long-term average relationships (except MEve). As chlorophyll *a* does not capture variation in phytoplankton composition or stoichiometric quality, these results further suggest that while biomass responds directly to resource availability, taxonomic structure is more strongly shaped by resource quality and niche differentiation

Nitrogen concentrations influenced short-term diversity dynamics but had no effect on long-term averages (except for concentration and MRic), highlighting that context-dependent, sometimes positive or negative effects shape zooplankton diversity temporal patterns without producing consistent mean changes. This, together with best predictability at intermediate to long time lags (3-4 weeks) matches their influence via indirect pathways such as phytoplankton quantity or quality. Contrary to its known importance for certain taxa and influence on lake zooplankton community structure (Shurin et al. 2010, Li et al. 2022), PO_4_ and SiOh4 had surprisingly little effect in our system. PO_4_ only was associated with changes in morphological evenness - negative on the long-term, but predominantly positive on the short-term dynamics - while SiOH_4_ was only relevant for MRic dynamics. This highlights their importance for few taxa without shaping overall community structure or concentration.

In contrast, particulate organic matter influenced long-term patterns of community structure, but played a minor role as regulator of dynamics, suggesting its association is largely correlative rather than mechanistic. Indeed, PON and POC reflect trophic state, phytoplankton or detritus biomass, and therefore likely co-vary with zooplankton due to shared seasonal cycles and long-term trends. Both compounds have declined significantly over the past 12 years at our study site (Beck et al. 2023), which may explain the apparent long-term associations. Matching this claim, PON and POC were identified as causal predictors of short-term dynamics only at immediate or short time lags (0/1 week).

Finally, results regarding salinity confirm its role as predictor of offshore water masses at Point B (but also mixing). It is part of the zooplankton dynamic system, but only with low/moderate predictive skill and low average effect identified in SMap. Its negative association in the long-term suggests that more offshore conditions reduce plankton concentration. Its identification as dynamic driver, notably at immediate or 1-week lags, further confirms its mechanistic link to water mass dynamics. The driver effects on zooplankton dynamics do not vary across years (S5), suggesting system stability despite significant long-term trends in most environmental variables (Beck et al. 2023).

Our analysis further provides insights into the nonlinearity of plankton dynamics which is crucial because the effects of environmental changes on communities can vary dramatically depending on the system’s state, in which case average responses may fail to capture key ecological processes. Further, our analyses revealed marked differences in nonlinearity among diversity metrics. Zooplankton concentration and morphological diversity exhibited moderate state dependence suggesting relatively linear responses to environmental forcing (and continuous, more or less variable, demographic processes). In contrast, taxonomic diversity exhibited higher predictability despite its stronger nonlinearity, indicating a closer, yet context-dependent, coupling to environmental conditions. This implies that while abundance and morphological trait-space structure respond gradually to environmental conditions, species identities reorganize in a more abrupt, context-dependent way.

Overall, zooplankton concentration and, notably morphological diversity, were less predictable than taxonomic diversity, although the predictors in the models captured a substantial part of the system’s dynamics (Sugihara et al. 2012). In the case of morphological diversity, this pattern likely reflects the aggregation of species into a reduced trait space, which can dampen species-specific responses and smooth variability, thereby weakening the apparent coupling with environmental forcing. Functional redundancy among taxa may further contribute to this pattern. Nevertheless, some variation still arises from unmeasured factors or stochastic events.

The divergence between long-term associations and dynamically inferred drivers has implications for predicting ecosystem responses to environmental change. Notably, our results highlight that conventional approaches may misidentify key environmental controls in nonlinear, state-dependent systems: GLMs would identify particulate organic matters as important drivers, but insights from EDM shows that they are likely largely correlative rather than mechanistic. Similarly, average effect approaches would have missed the dynamic signal of morphological homogenisation or underestimated the possible impact of warming on the zooplankton community. Predictions of ecosystem responses to environmental change based solely on average relationships may be incomplete or misleading.

The more linear responses of morphological diversity and zooplankton concentration indicate that linear approaches could capture their broad patterns reasonably well, even if fine-scale, short-term dynamics remain less predictable, whereas linear models might be less appropriate for predicting dynamics in taxonomic diversity.

Methodologically, the two analytical approaches relied on differently pre-processes input data (de-trended for CCM/SMap, de-seasonalised for GLM) to satisfy model requirements and ensure unbiased ecological interpretation of estimated effects. This prevents direct comparison of model performance, which was not this study’s objective. In addition, CCM itself does not distinguish between direct and indirect drivers. However, only variables whose fluctuations propagate through the system and systematically influence a diversity variable are identified as causal; purely correlated but dynamically irrelevant variation would not be detected. Although each SMap fit is locally linear, the coefficients vary across reconstructed states, allowing the model to capture nonlinear, state-dependent interactions. In this sense, SMap provides a dynamic view of ecological drivers and reveals how the influence of each predictor changes over time.

While the 12-year time series captures substantial interannual variability, it may not fully resolve slower, multi-decadal processes or regime shifts, and thus primarily reflects medium-term community–environment dynamics.

We note that some of the weak relationships we found regarding the zooplankton dynamics – being identified ‘causal’ but showing low effect strong across time – might indicate their very local importance for certain taxa, but with only minor effect on community structure.

## Conclusion

Our study demonstrates that zooplankton diversity is shaped by both continuous and context-dependent environmental drivers. Temperature emerged as a continuous, pervasive driver, whereas nutrients and phytoplankton mainly acted through shorter-term or indirect pathways. Importantly, we find evidence for morphological homogenisation in response to short-term fluctuations in chlorophyll *a*,which was not detectable in long-term average relationships. By combining nonlinear, state-dependent analyses with estimation of long-term average effects, we show that the identity and influence of environmental drivers can vary across temporal scales and diversity dimensions, highlighting that mean environmental associations do not necessarily reflect the mechanisms governing temporal change in zooplankton communities.

## Data availability

Time series data is available in Beck et al. (2023). All code used to conduct the analysis is accessible on github/codeberg.

## Acknowledgements

Thanks to CESAB and IMPACTS group for the support.

