## Supplementary material for "Mean environmental associations obscure drivers of zooplankton community dynamics"

*S1: Characteristics of pairwise relationships between environmental variables and diversity dynamics. For each environmental variable, only the time lag resulting in highest prediction skill ('rho') is provided. Bold writing indicates causal drivers beyond seasonal signal, as identified from Kendall tau, seasonal surrogates and t.test.*

| Div | Env | Rho | E_opt | Lag_opt | kendall.p | kendall.tau | surr_avg.p | diffzero.p |
|---|---|---|---|---|---|---|---|---|
| Conc | **Chl *a*** | **0.4141** | **3** | **-2** | **<0.0001** | **0.65977** | **0.0099** | **<0.0001** |
| | **Temperature** | **0.5019** | **3** | **-3** | **<0.0001** | **0.95862** | **0.0099** | **<0.0001** |
| | **Salinity** | **0.2376** | **3** | **0** | **<0.0001** | **0.76092** | **0.0099** | **<0.0001** |
| | **Oxygen** | **0.2141** | **3** | **0** | **<0.0001** | **0.8023** | **0.0099** | **<0.0001** |
| | NO2 | 0.3649 | 3 | -5 | 0.6514 | -0.04828 | NA | <0.0001 |
| | NO3 | 0.3285 | 3 | -5 | 0.9986 | -0.37471 | NA | <0.0001 |
| | PO4 | 0.0892 | 3 | -1 | <0.0001 | 0.82529 | 0.1188 | <0.0001 |
| | **SiOH4** | **0.1301** | **3** | **-5** | **1.00E-04** | **0.45287** | **0.0396** | **<0.0001** |
| | **POC** | **0.1501** | **3** | **-2** | **<0.0001** | **0.55402** | **0.0099** | **<0.0001** |
| | **PON** | **0.2018** | **3** | **-2** | **<0.0001** | **0.50805** | **0.0099** | **<0.0001** |
| TRic | Chl *a* | 0.2504 | 3 | -5 | 0.5422 | -0.01149 | NA | <0.0001 |
| | **Temperature** | **0.5491** | **3** | **-3** | **0.0188** | **0.26897** | **0.0099** | **<0.0001** |
| | **Salinity** | **0.1605** | **3** | **-5** | **<0.0001** | **0.95402** | **0.0198** | **<0.0001** |
| | **Oxygen** | **0.2372** | **3** | **0** | **<0.0001** | **0.63218** | **0.0099** | **<0.0001** |
| | **NO2** | **0.3999** | **3** | **-5** | **<0.0001** | **0.56322** | **0.0099** | **<0.0001** |
| | **NO3** | **0.3621** | **3** | **-3** | **<0.0001** | **0.8023** | **0.0099** | **<0.0001** |
| | **PO4** | **0.1007** | **3** | **0** | **<0.0001** | **0.90345** | **0.0198** | **<0.0001** |
| | SiOH4 | 0.2074 | 3 | -3 | 1 | -0.63678 | NA | <0.0001 |
| | **POC** | **0.0822** | **3** | **0** | **<0.0001** | **0.84828** | **0.0495** | **<0.0001** |
| | **PON** | **0.1164** | **3** | **-2** | **<0.0001** | **0.85287** | **0.0099** | **<0.0001** |
| TShannon | **Chl *a*** | **0.5367** | **3** | **-1** | **<0.0001** | **0.90805** | **0.0099** | **<0.0001** |
| | **Temperature** | **0.7402** | **3** | **-1** | **<0.0001** | **0.83448** | **0.0099** | **<0.0001** |
| | **Salinity** | **0.237** | **3** | **-2** | **<0.0001** | **0.8023** | **0.0099** | **<0.0001** |
| | **Oxygen** | **0.3084** | **3** | **-2** | **<0.0001** | **0.92184** | **0.0099** | **<0.0001** |
| | **NO2** | **0.5888** | **3** | **-3** | **<0.0001** | **0.89885** | **0.0099** | **<0.0001** |
| | **NO3** | **0.5399** | **3** | **-5** | **<0.0001** | **0.69195** | **0.0099** | **<0.0001** |
| | **PO4** | **0.1015** | **3** | **-4** | **<0.0001** | **0.67356** | **0.0495** | **<0.0001** |
| | **SiOH4** | **0.3478** | **3** | **-1** | **<0.0001** | **0.86667** | **0.0099** | **<0.0001** |
| | **POC** | **0.1203** | **3** | **-1** | **<0.0001** | **0.84368** | **0.0099** | **<0.0001** |
| | **PON** | **0.2765** | **3** | **0** | **<0.0001** | **0.72414** | **0.0099** | **<0.0001** |
| TPielou | Chl *a* | 0.4847 | 3 | -2 | 0.6772 | -0.05747 | NA | <0.0001 |
| | **Temperature** | **0.7341** | **3** | **-3** | **9.00E-04** | **0.3977** | **0.0099** | **<0.0001** |
| | Salinity | 0.187 | 3 | 0 | 0.8794 | -0.14943 | NA | <0.0001 |
| | **Oxygen** | **0.304** | **3** | **-2** | **<0.0001** | **0.95402** | **0.0099** | **<0.0001** |
| | NO2 | 0.5941 | 3 | -5 | 1 | -0.55402 | NA | <0.0001 |
| | NO3 | 0.5048 | 3 | -5 | 0.8864 | -0.15402 | NA | <0.0001 |
| | PO4 | 0.0715 | 3 | 0 | 0.4718 | 0.01149 | NA | <0.0001 |
| | **SiOH4** | **0.3674** | **3** | **-1** | **<0.0001** | **0.91264** | **0.0099** | **<0.0001** |
| | **POC** | **0.1536** | **3** | **-1** | **<0.0001** | **0.84368** | **0.0099** | **<0.0001** |
| | **PON** | **0.2676** | **3** | **0** | **0.0245** | **0.25517** | **0.0099** | **<0.0001** |
| MRic | **Chl *a*** | **0.2772** | **4** | **-2** | **<0.0001** | **0.78391** | **0.0099** | **<0.0001** |
| | Temperature | 0.4625 | 4 | -4 | 0.9843 | -0.27356 | NA | <0.0001 |
| | Salinity | -0.04 | 4 | -2 | 1 | -0.63678 | NA | 1 |

| | | | | | | | | |
|---|---|---|---|---|---|---|---|---|
| | Oxygen | **0.1685** | **4** | **-1** | **<0.0001** | **0.65057** | **0.0099** | **<0.0001** |
| | NO2 | 0.2576 | 4 | -5 | 1 | -0.72414 | NA | <0.0001 |
| | NO3 | 0.3187 | 4 | -5 | 0.1783 | 0.12184 | NA | <0.0001 |
| | PO4 | -3.00E-04 | 4 | -5 | 0.5978 | -0.02989 | NA | 0.7478 |
| | SiOH4 | **0.2408** | **4** | **-4** | **<0.0001** | **0.82069** | **0.0099** | **<0.0001** |
| | POC | **0.1215** | **4** | **-2** | **<0.0001** | **0.92644** | **0.0198** | **<0.0001** |
| | PON | 0.0693 | 4 | -2 | 0.3102 | 0.06667 | NA | <0.0001 |
| MDiv | **Chl *a*** | **0.2816** | **3** | **-4** | **<0.0001** | **0.82529** | **0.0099** | **<0.0001** |
| | Temperature | 0.3775 | 3 | -4 | 0.8308 | -0.12184 | NA | <0.0001 |
| | Salinity | **0.156** | **3** | **0** | **5.00E-04** | **0.41609** | **0.0099** | **<0.0001** |
| | Oxygen | **0.1184** | **3** | **-5** | **2.00E-04** | **0.44368** | **0.0297** | **<0.0001** |
| | NO2 | **0.2693** | **3** | **-5** | **0.0107** | **0.29655** | **0.0099** | **<0.0001** |
| | NO3 | **0.2652** | **3** | **-4** | **0.0064** | **0.31954** | **0.0099** | **<0.0001** |
| | PO4 | **0.2551** | **3** | **-4** | **<0.0001** | **0.91724** | **0.0099** | **<0.0001** |
| | SiOH4 | 0.1348 | 3 | -2 | 0.9812 | -0.26437 | NA | <0.0001 |
| | POC | **0.1327** | **3** | **0** | **<0.0001** | **0.88966** | **0.0198** | **<0.0001** |
| | PON | **0.2049** | **3** | **0** | **<0.0001** | **0.75172** | **0.0099** | **<0.0001** |
| MEve | Chl *a* | 0.417 | 5 | -4 | 0.2076 | 0.10805 | NA | <0.0001 |
| | Temperature | 0.4808 | 5 | -4 | 0.262 | 0.08506 | NA | <0.0001 |
| | Salinity | **0.1488** | **5** | **0** | **<0.0001** | **0.48966** | **0.0099** | **<0.0001** |
| | Oxygen | **0.2135** | **5** | **0** | **<0.0001** | **0.88506** | **0.0099** | **<0.0001** |
| | NO2 | 0.4115 | 5 | -5 | 0.9709 | -0.24138 | NA | <0.0001 |
| | NO3 | 0.3422 | 5 | -5 | 0.9928 | -0.31034 | NA | <0.0001 |
| | PO4 | **0.2709** | **5** | **-5** | **<0.0001** | **0.93103** | **0.0099** | **<0.0001** |
| | SiOH4 | **0.2268** | **5** | **-5** | **<0.0001** | **0.91724** | **0.0099** | **<0.0001** |
| | POC | **0.1885** | **5** | **0** | **<0.0001** | **0.76552** | **0.0099** | **<0.0001** |
| | PON | **0.2598** | **5** | **0** | **<0.0001** | **0.75172** | **0.0099** | **<0.0001** |

*S2: Predictive skills of environment variables on diversity dynamics across time lags of 0 to 5 weeks. Rho values are a measure of strength of predictability which can be correlated to but is no measure of effect strength.*

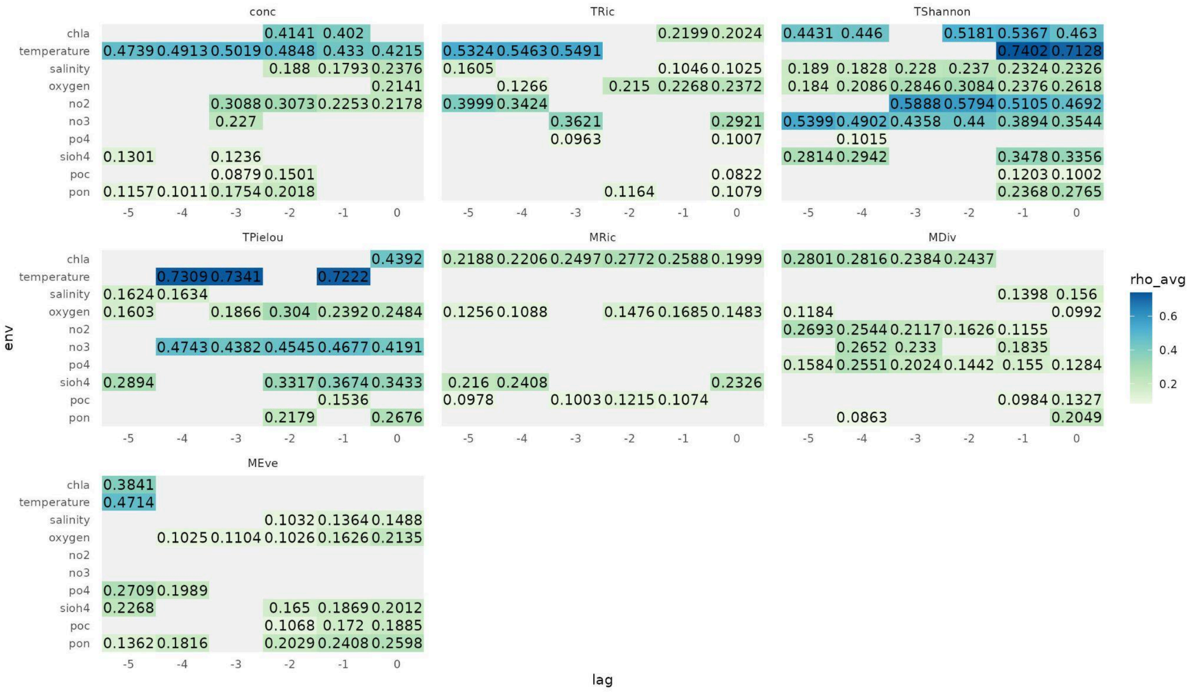


*S3: Model statistics and summarised coefficients of multivariate SMaps using the optimal time lag of each included predictor. Model rho value is a measure of model fit, theta indicates linearity of the system and E_opt the embedding used. E-1 predictors have been included in each model, selected with decreasing prediction skill in CCM. SMap coefficients for each predictor were summarised across the whole 12-year time series, to indicate mean effect and absolute mean effect on diversity, effect variability (SD) and % of positive values (%per).*

| Div | rho | Theta/ θ | E_opt | Env (-timelag) | Mean | $Mean_{abs}$ | SD | %pos |
|---|---|---|---|---|---|---|---|---|
| Conc | 0.4604 | 1 | 3 | Chl *a* (3) | 0.2541 | 0.2541 | 0.0811 | 1 |
| | 0.4604 | 1 | 3 | Temperature (2) | -0.2201 | 0.2201 | 0.0794 | 0 |
| TRic | 0.6083 | 4 | 3 | Temperature (3) | 0.6963 | 0.7798 | 0.6583 | 0.7319 |
| | 0.6083 | 4 | 3 | NO2 (5) | 0.0115 | 0.2793 | 0.1865 | 0.4772 |
| TShannon | 0.7831 | 3 | 3 | Temperature (1) | 0.5912 | 0.686 | 0.4748 | 0.7083 |
| | 0.7831 | 3 | 3 | NO2 (3) | -0.0914 | 0.2977 | 0.1895 | 0.3997 |
| TPielou | 0.7633 | 3 | 3 | Temperature (3) | 0.6224 | 0.6723 | 0.4761 | 0.7403 |
| | 0.7633 | 3 | 3 | NO3 (4) | -0.0713 | 0.2049 | 0.1375 | 0.4384 |
| MRic | 0.36 | 1,5 | 3 | Chl *a* (2) | -0.3389 | 0.3389 | 0.1042 | 0 |
| | 0.36 | 1,5 | 3 | SiOH4 (4) | -0.275 | 0.275 | 0.084 | 0 |
| MDiv | 0.3232 | 0.75 | 3 | Chl *a* (4) | -0.3158 | 0.3158 | 0.0462 | 0 |
| | 0.3232 | 0.75 | 3 | NO2 (5) | -0.0117 | 0.0245 | 0.0269 | 0.5093 |
| MEve | 0.5906 | 3 | 3 | Chl *a* (5) | -0.1935 | 0.2214 | 0.1463 | 0.1574 |
| | 0.5906 | 3 | 3 | Temperature (5) | 0.3679 | 0.378 | 0.2377 | 0.9323 |
| MEve | 0.5906 | 3 | 3 | PO4 (5) | 0.0645 | 0.1416 | 0.1029 | 0.6667 |

| | | | | | | | |
|---|---|---|---|---|---|---|---|
| 0.5906 | 3 | 3 | PON (0) | -0.0182 | 0.1342 | 0.099 | 0.4484 |

*S4: Average absolute effect strength ('abs-mean', A), effect variability ('SD', B) and proportion of positive effects ( C) of environmental drivers on diversity. SMap coefficients were obtained from multivariate SMAP using all predictors identified as causal drivers in CCM at the time lag of their highest prediction skill. The embedding dimension was set to $n_{pred}$+1. Using suboptimal E will result in less accurate estimators (Sugihara et al. 2012), but can be a practical solution to describe essential behaviour of the attractor (Wang et al. 2020)]. SMap coefficients for each predictor were summarised across the whole 12-year time series.*

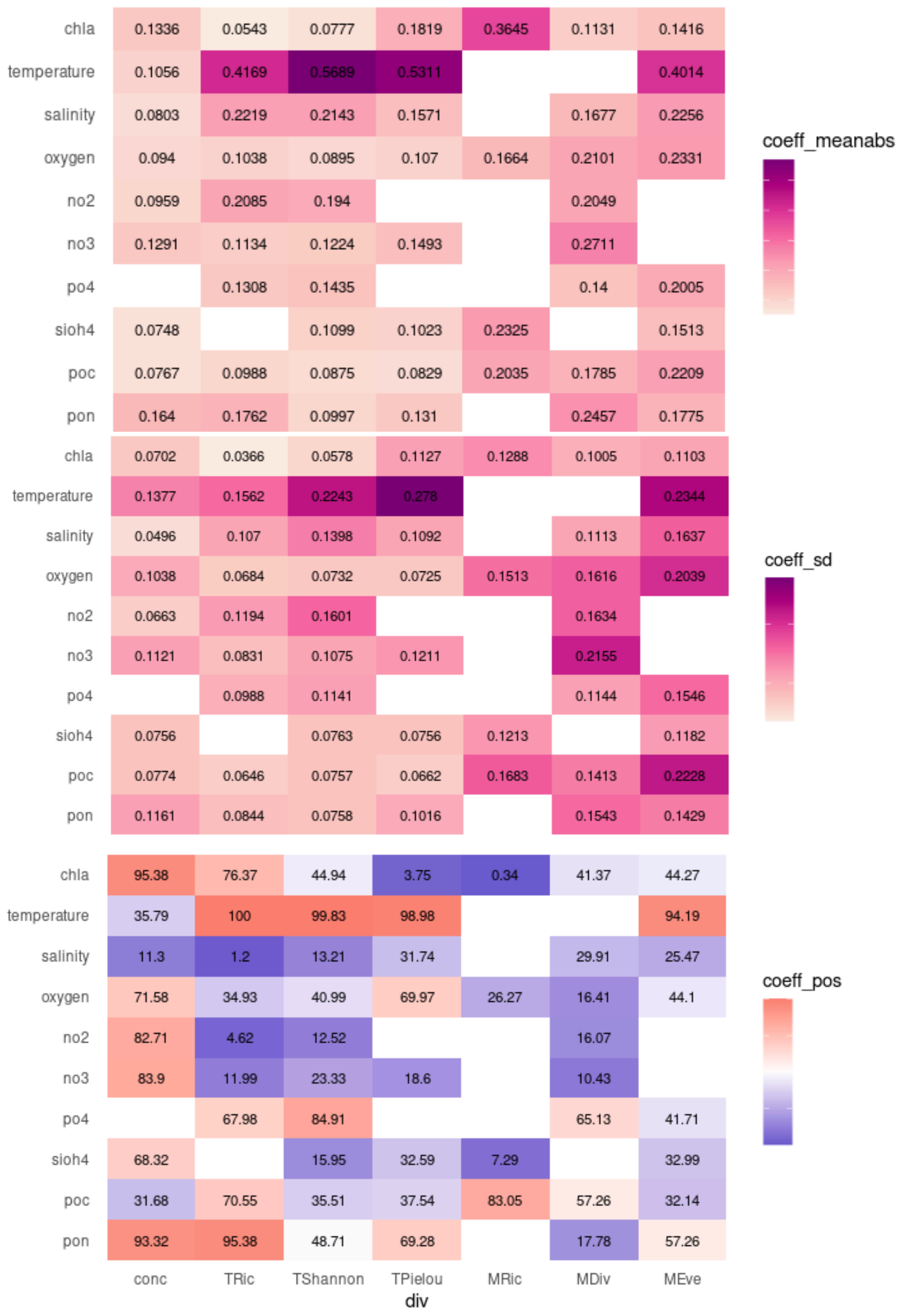

*S5: SMap coefficient across the study period, time indicates timestep*

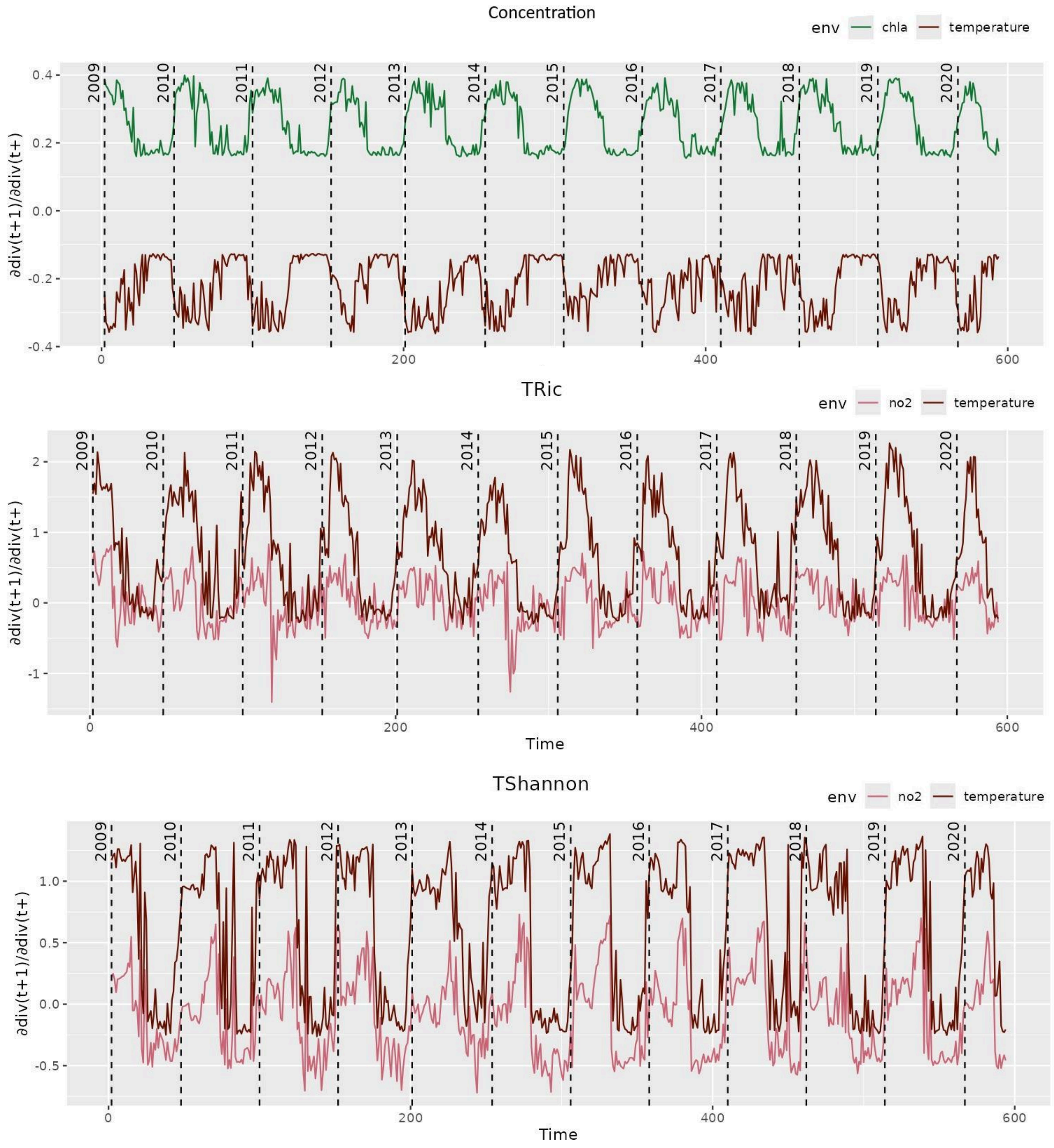

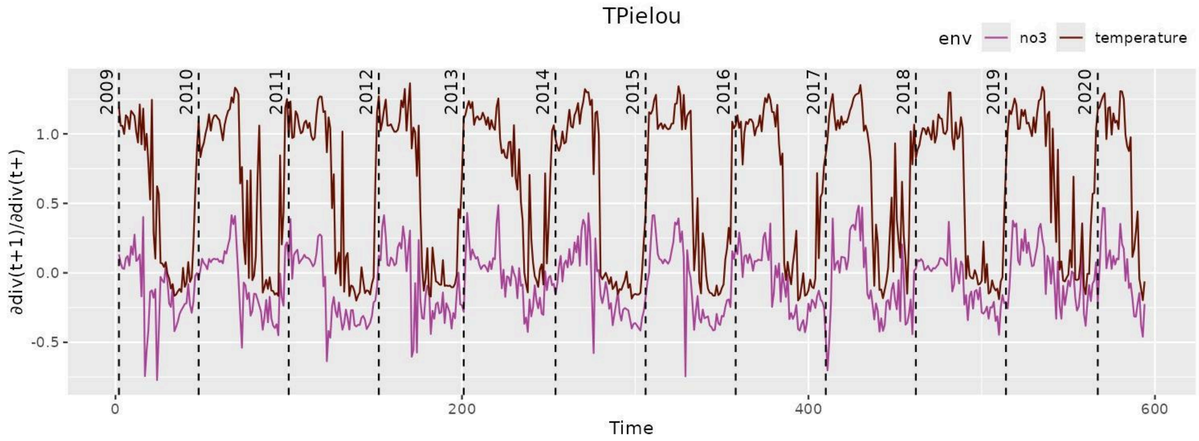
TPielou
env
no3
temperature
∂div(t+1)/∂div(t+)
Time
2009
2010
2011
2012
2013
2014
2015
2016
2017
2018
2019
2020
1.0
0.5
0.0
-0.5
0
200
400
600

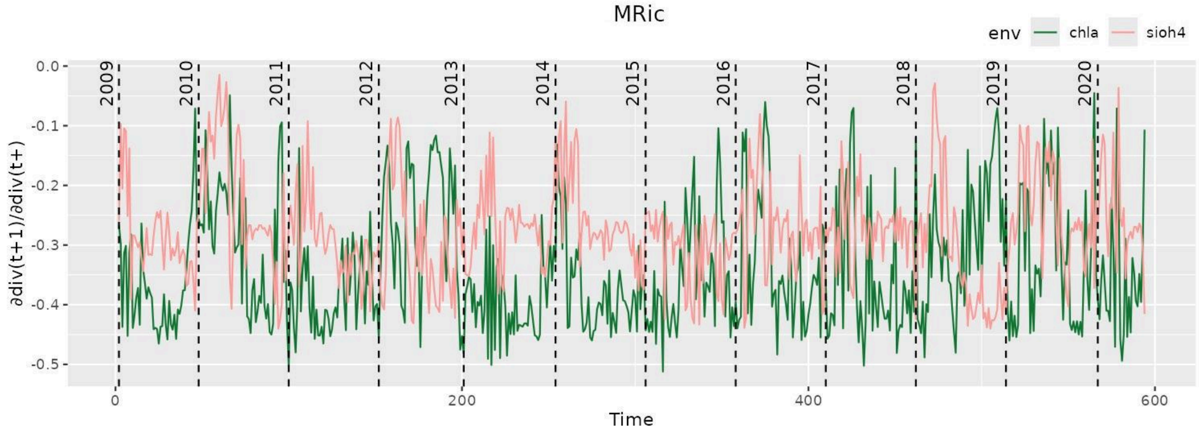
MRic
env
chla
sioh4
∂div(t+1)/∂div(t+)
Time
2009
2010
2011
2012
2013
2014
2015
2016
2017
2018
2019
2020
0.0
-0.1
-0.2
-0.3
-0.4
-0.5
0
200
400
600

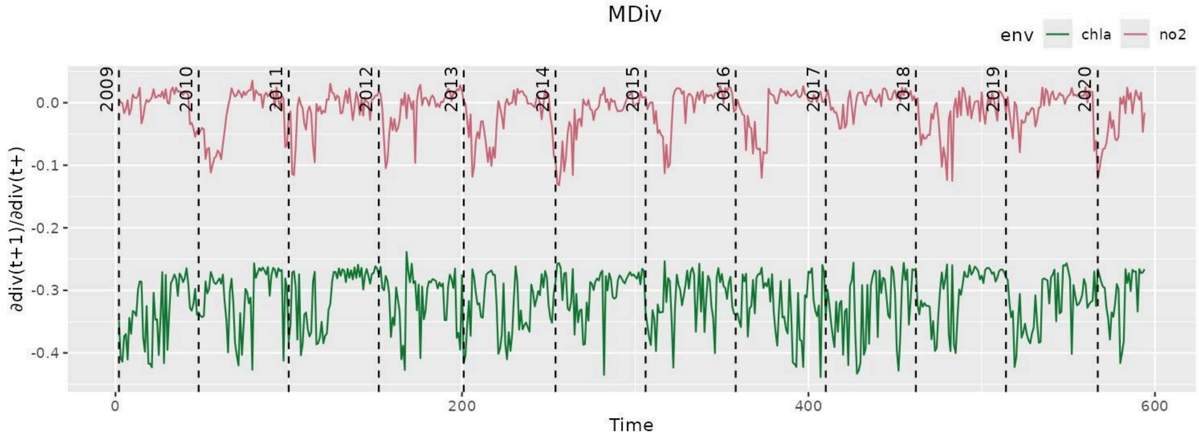
MDiv
env
chla
no2
∂div(t+1)/∂div(t+)
Time
2009
2010
2011
2012
2013
2014
2015
2016
2017
2018
2019
2020
0.0
-0.1
-0.2
-0.3
-0.4
0
200
400
600

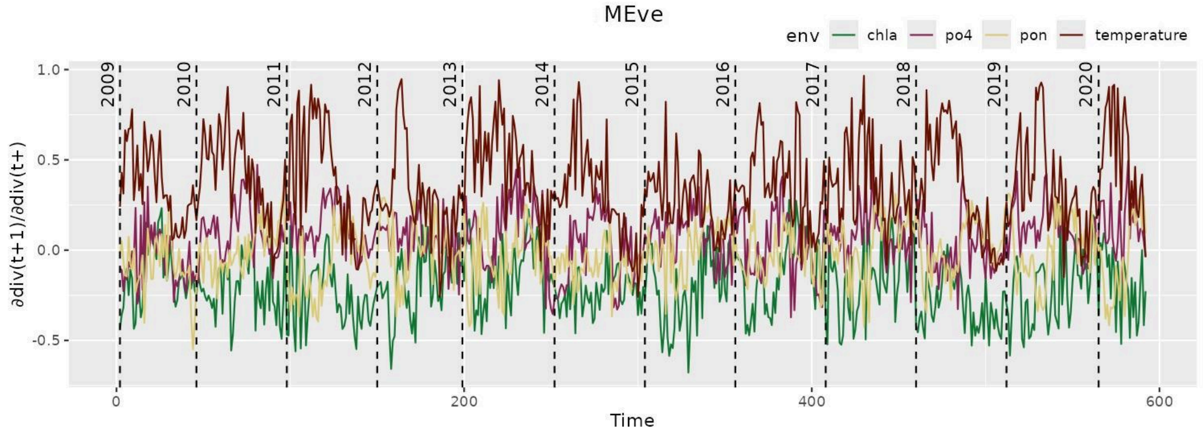
MEve
env
chla
po4
pon
temperature
∂div(t+1)/∂div(t+)
Time
2009
2010
2011
2012
2013
2014
2015
2016
2017
2018
2019
2020
1.0
0.5
0.0
-0.5
0
200
400
600

*S6: Yearly averaged SMap coefficient (mean) across the study period.*

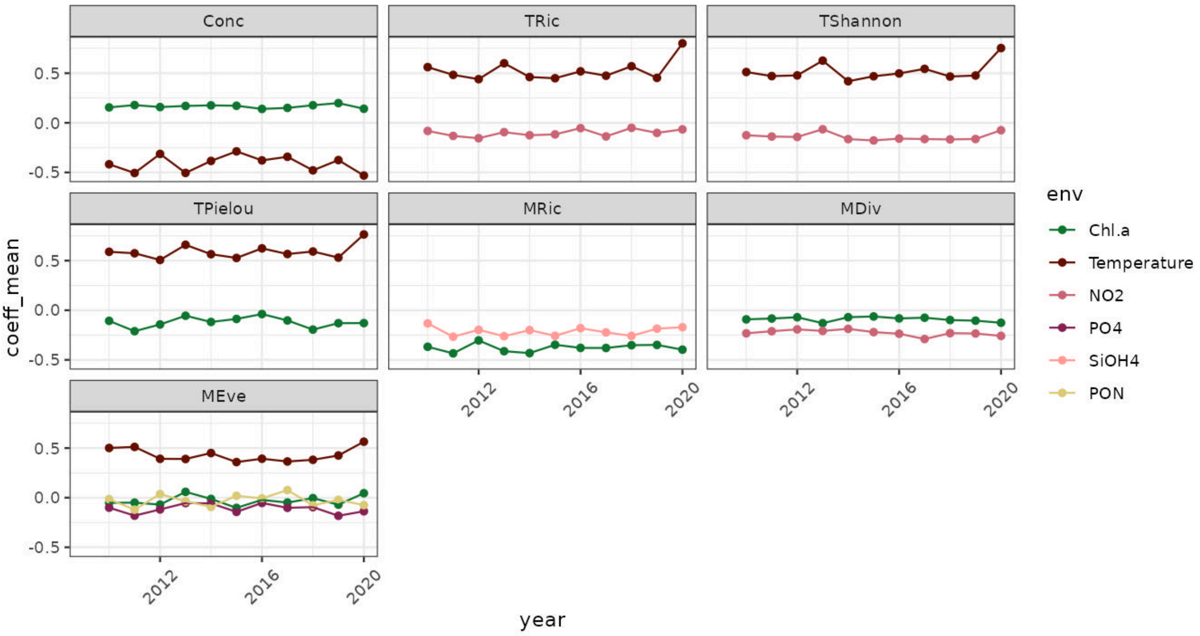


*S7: Predictive skills of raw or aggregated environment variables on diversity dynamics across time lags of 0 to 5 weeks. Aggregated environmental variables are average values over two or three weeks. Color reflects rho values, which are a measure of strength of predictability which can be correlated to but is no measure of effect strength.*

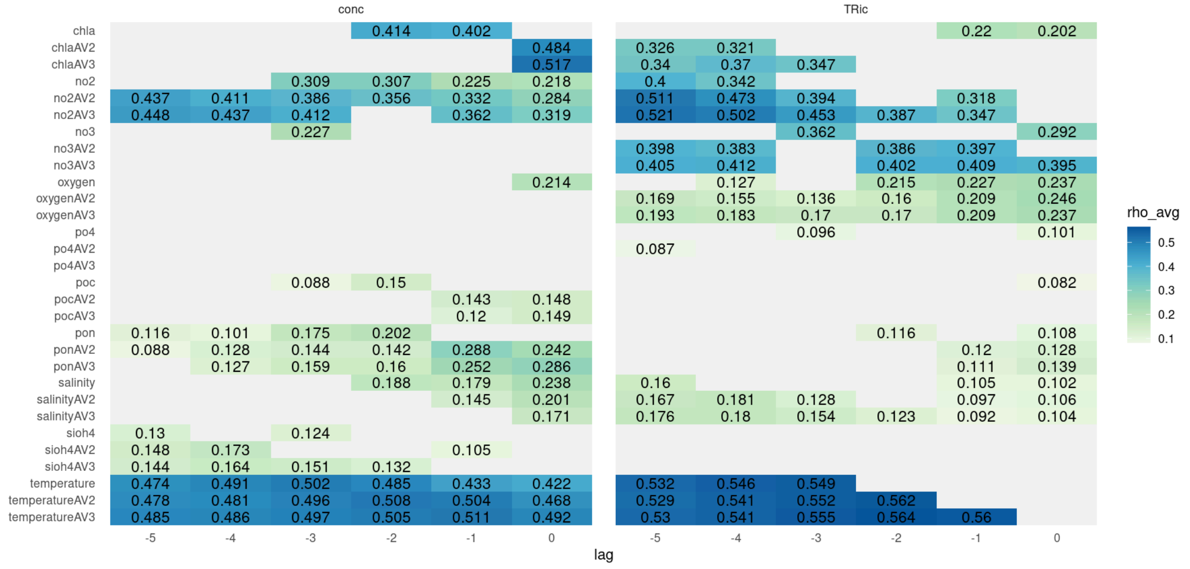
